# The biochemical properties of a novel paraoxonase-like enzyme in *Trichoderma atroviride* strain T23 involved in the degradation of 2,2-dichlorovinyl dimethyl phosphate

**DOI:** 10.1101/416081

**Authors:** Jianan Sun, Xu Yuan, Yaqian Li, Xinhua Wang, Jie Chen

## Abstract

Dichlorvos, is a broad-spectrum organophosphorus pesticide that is widely applied in the agricultural industry and considered a pollutant to fish and bees. *T. atroviride* strain T23, an efficient DDVP-degrading strain, could convert DDVP to dichloroacetic acid, 2,2-dichloroethanol and phosphoric acid through mineralization. RT-qPCR analysis showed *TaPon1*-like encoding an organophosphorus hydrolase, is continuously highly expressed in the process of degrading DDVP. *TaPon1-*like contained an open reading frame of 1317 bp, and the deduced amino acid sequence shared 21% homology with HuPON1, which also exhibits excellent hydrolysis of organophosphate-oxons compounds. By analysis of gene knockout, we found the Δ*TaPon1-*like knockout strain KO1 lost 35.6% of its DDVP-degradation capacity at 24 h, but this loss of degradation activity was recovered when the gene was complemented. Furthermore, the purified recombinant protein reTAPON1-LIKE, could transform DDVP only to dimethyl phosphate and showed significant paraoxonase activity (1028 U L^−1^). The reTAPON1-LIKE enzyme showed a broad degradation spectrum, degrading not only DDVP but also organophosphate-oxons and lactone. The kinetic parameters (*Km* and *kcat*) of the purified reTAPON1-LIKE were determined to be 0.23 mM and 204.3 s^−1^ for DDVP, respectively. The highest activity was obtained at 35 °C, and the optimal pH was 8.5. The activity of reTAPON1-LIKE was enhanced most significantly when 1.0 mM Ca^2+^ was added but declined when 1.0 mM Cu^2+^ was added. These results showed TAPON1-LIKE play an important role for DDVP degradation in the first step by T23 and provided clue to comprehensively understanding the degradation mechanism of organophosphate-oxons pesticides by filamentous fungi.

**Importance:** The large amounts of residues of organophosphate pesticides in agroecological system has become a great threat to the safety of environment and humans. Bioremediation in association with microbial is innovative technology having a potential to alleviate such pollution problems. The genus *Trichoderma* is genetically diverse with capabilities to degrade chemical pesticides among different strains with agricultural significance. As a typical organophosphorus pesticide, it is one of the most employed compounds of the family. Though it was classified as a highly toxic pesticide by WHO due to its hazardous properties, it plays an important role in the control of plant pests, food storage and homes, as well as to treat infections in livestock. Therefore, we use DDVP as a model of organophosphate pesticide to study the mechanism of *Trichoderma* degrading organophosphate pesticides, for the aim of globally understanding molecular mechanism of enzymatic degradation of organophosphate pesticides by beneficial fungi.

## Introduction

Organophosphorus pesticides are some of the most widely used pesticides. Dichlorvos, 2,2-dichlorovinyl dimethyl phosphate (DDVP), is a broad-spectrum organophosphorus pesticide often applied to agricultural crops and forests and in aquatic environments (1). DDVP, although usually viewed as a moderately toxic pesticide, is a pollutant to fishes and bees. In China, the demand for DDVP was estimated to be over 40,000 tons in 2007, and its usage is expected to increase since five highly toxic organophosphate insecticides (e.g., parathion) have been banned (2). However, severe contamination may arise due to the widespread use and discharge of DDVP into the environment, and its residues can be detected in water, soil, vegetables, fruits, milk, and living organisms (3). In addition, DDVP is highly toxic to nontarget invertebrates and vertebrates, resulting in irreversible inhibition of acetylcholinesterase, which is needed for regulating the neurotransmitter acetylcholine (4). Therefore, developing a high-efficiency method to remove DDVP residues is necessary for environmental and food production safety.

Conventional methods for the removal of DDVP, such as chemical hydrolysis/oxidation (2) and photocatalytic oxidation (5), are difficult to apply to large contaminated areas and are very expensive. Biodegradation depending on microbial metabolism has become an attractive approach for removing hazardous chemicals such as organophosphate pesticides from the environment (6). Moreover, some selected microbes have the capacity to degrade DDVP (7), and these strains can usually be isolated from natural environments.

The genus *Trichoderma* has attained a unique position in the agricultural industry as a successful biocontrol agent against plant diseases, plant growth promoters and soil bioremediation (8). More importantly, *Trichoderma* spp. with multiple functions in soil remediation, such as removing heavy metals and chemical pesticide residues, particularly when applied together with hyperaccumulators, have been revealed. Zhang et al. (9) isolated and characterized the *Trichoderma* strain TC5, which has high degradation activity against chlorpyrifos. In previous work, we reported that *T. atroviride* strain T23 has the capacity to degrade DDVP and that the key genes *hex1* (10) and *Tapdr2* (11) were associated with the tolerance of *T. atroviride* strain T23 to DDVP.

Additionally, several genes involved in the degradation of organophosphorus compounds, including *opd, opa, opdA*, and *mpd*, have been discovered and cloned from different bacterial species (12). However, there are few studies on organophosphorus pesticide degradation enzymes from fungi; for instance, the crucial enzymes and related genes responsible for organophosphate pesticide degradation in strain T23 have not been reported.

Luckily, some molecular details of the DDVP metabolic pathway in humans is known. For instance, paraoxonases from humans (HuPONs) have been confirmed to function in the degradation of DDVP (13). It was supposed that these genes might also be present in *T. atroviride*. Therefore, the function of these genes in the biodegradation of organophosphorus pesticides needs to be investigated. This study aimed to demonstrate whether *HuPon*s were present in *T. atroviride* strain T23 and what function strain T23 plays in the biodegradation of DDVP.

## Results

The effects of DDVP on the growth of T23 and biodegradation

In Fig. 1A, DDVP degradation by strain T23 was observed after DDVP addition at 24 h. DDVP may inhibit the growth of train T23, and especially high concentrations of DDVP (≥ 400 μg mL^−1^) were toxic to fungal growth and inhibited degradation. While DDVP existed at 100-300 μg mL^−1^ concentration, there was a weak decrease in biomass of strain T23. Even at DDVP concentrations up to 300 μg mL^−1^, the degradation rate reached 56.7%, which is more than the DDVP self-degradation rate. Strain T23 grew rapidly when DDVP was inoculated in the first 24 hours and the growth curve of strain with or without DDVP showed a similar trend (Fig. 1B).

**Fig. 1.**
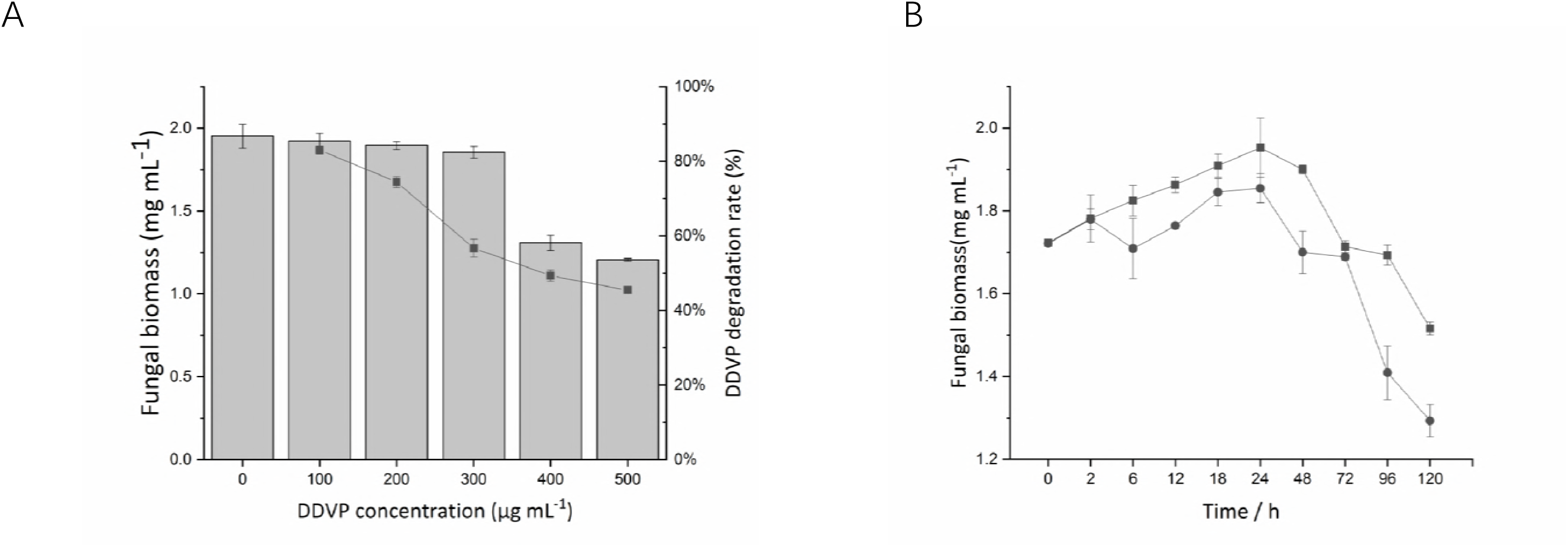
Fungal growth and DDVP degradation rate in cultures of *T. atroviride* T23. (A) Effects of DDVP concentration on fungal biomass (column) and DDVP degradation rate (solid square). DDVP was added at the beginning of the cultures. The biomass and DDVP degradation were determined after 24 h of incubation. (B) Effect of incubation time on fungal biomass. DDVP at 300 μg mL^−1^ was added at the beginning of the cultures. The group without DDVP was used as a control. Solid square, control; solid circle, under 300 μg mL^−1^ DDVP. Each value is expressed as the means ± standard errors of three replicates.

An assessment of morphological changes in response to DDVP (300 μg mL^−1^) accumulation in strain T23 and the quantification of DDVP were performed by SEM-EDS analysis. SEM analysis of mycelia was performed at 6, 24, and 72 h of incubation. No peak of chlorine was detected by EDS (Fig. S1) and the adsorbate concentration of DDVP detected using a GC-FPD showed no peak (data not shown), ether. According to these results, degradation of DDVP by mycelial adsorption was excluded and the enzymes that are strain T23 products are the primary factor attributed to DDVP degradation.

DDVP metabolite identification

To clarify the DDVP degradation pathway of strain T23, the intermediate metabolites produced during DDVP degradation were analyzed and identified by GC-MS. DDVP with a retention time (RT) of 13.774 min (Fig. 2A) was still present in the medium after five days. By comparing the extractions following derivatization of degraded and non-degraded DDVP, GC-MS analysis of the samples gave three significantly different peaks with RTs of 13.075, 14.115, and 21.83 min (Fig. 2A) representing metabolites *tert*-butyldimethylsilyl derivative of 2,2-dichloroethanol, dimethyl phosphate, and phosphoric acid, respectively. By comparing the anhydrous ethyl ether extractions of degraded and non-degraded DDVP, GC-MS analysis of the samples gave two significantly different peaks with RTs of 5.577 and 10.606 min (Fig. 2B) representing metabolites of 2,2-dichloroethanol and phosphoric acid.

**Fig. 2.**
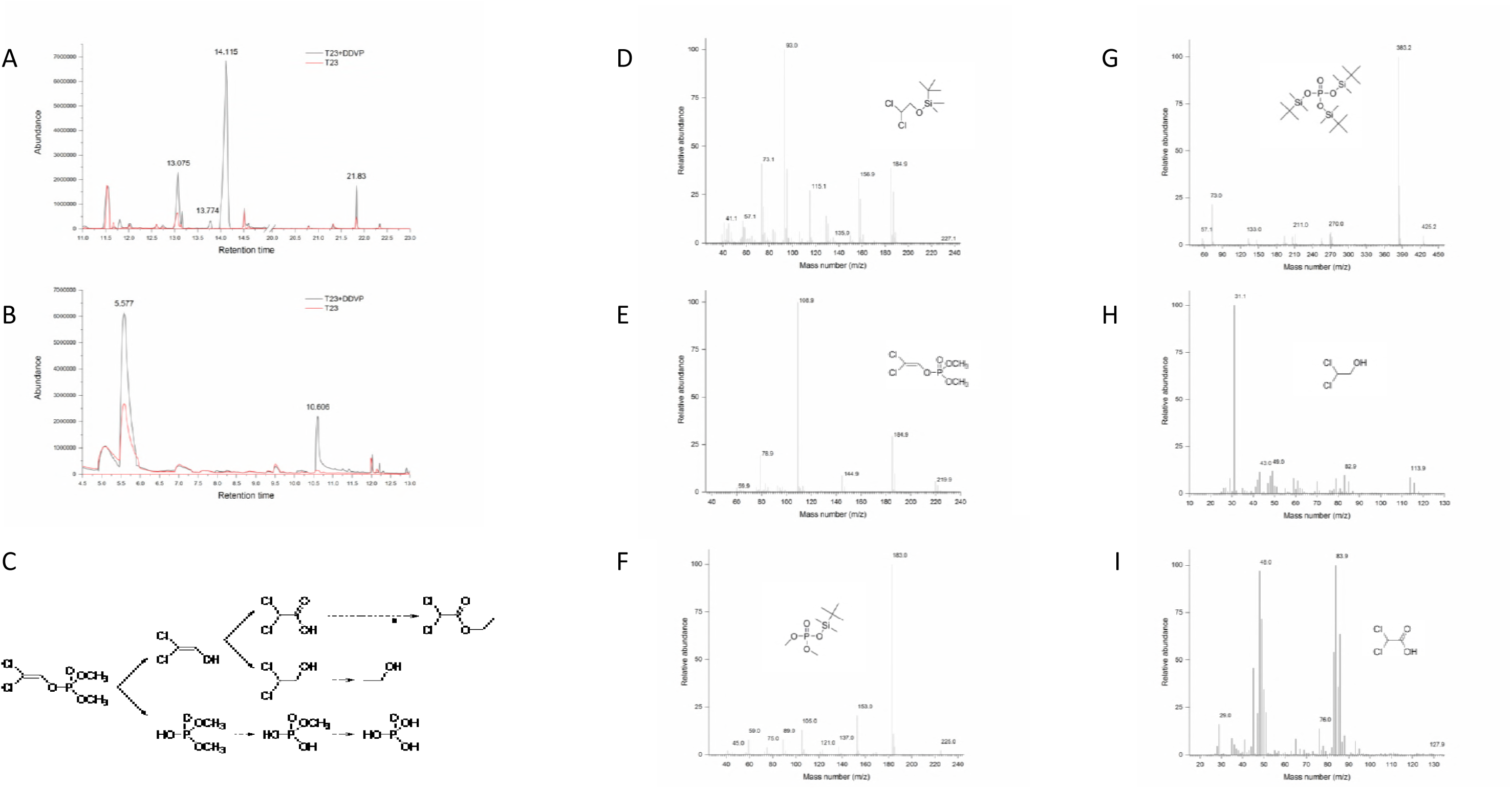
Gas chromatogram and mass spectrum detection of DDVP degradation by strain T23. A gas chromatogram of the extract obtained from the culture at 5 days treated with the MTBSTFA (A) and non-treated (B). The black line indicated strain T23 incubate in Burk medium with initial concentration of 300 μg mL^−1^ DDVP and the red line indicated strain T23 incubate in Burk medium. Potential catabolic pathway for DDVP degradation by strain T23 (C). Mass spectra of DDVP (E) degradation products formed the *tert*-butyldimethylsilyl derivatives were identified of the peak with RTs of 13.075 min (D), 14.115 min (F) and 21.83min (G). Mass spectra of DDVP (E) degradation products were identified of the peak with RTs of 5.577 min (H) and 10.606min (I).

Each of the six peaks was identified according to its mass spectrum and the NIST library identi?cation program. Compound E had the same retention time as the DDVP standard (RT = 13.774 min). The mass spectral data also demonstrated that compound E was DDVP (Fig. 2E). 2,2-dichloroethanol was identified following derivatization, which yielded the *tert*-Butyldimethylsilyl derivative of 2,2-dichloroethanol [Cl2CHCH2OSi (CH3)2C(CH3)3, *m/z* 228]. The mass spectrum (Fig. 2D) of this molecule showed loss of H^+^ at *m/z* 227.1, and mass of fragments at *m/z* 156.9 [Cl2CHC(O)OSiH2], 115.1 [*tert*-Butyldimethylsilyl], and 93.0 [ClCH2C(O)O] were evident. The mass spectrum of 2,2-dichloroethanol (Fig. 2H) showed a prominent molecular ion at *m/z* 113.9 [M]^+^, and the fragments included *m/z* 82.9 [Cl2CH], 79 [ClCHCH2OH], 49 [ClCH2], 43 [C2H3O] and 31.1 [CH3O]. Chemical ionization and a caparison with two mass spectra (Fig. 2D & Fig. 2H) of authentic chemical confirmed the identity of 2,2-dichloroethanol. The mass spectrum of the *tert*-Butyldimethylsilyl derivative of dimethyl phosphate was shown in Fig. 2F. Although the molecular ion ([M]^+^, *m/z* 240) was not observed, the characteristic loss of *tert*-butyl (*m/z* 183) and a loss of methyl (m/ z 225) were noted. The molecular ion ([M]^+^, *m/z* 440) of the *tert*-Butyldimethylsilyl derivative of phosphoric acid (Fig. 2G) was not observed, but a characteristic loss of methyl (*m/z* 425.2) and a loss of *tert*-butyl (*m/z* 383.2) were confirmed. Compound H with RT of 10.606m showed a molecular ion at *m/z* 127.9 [M]^+^, which corresponds to dichloroacetic acid (Fig. 2H). The fragments included *m/z* at 83.9 [Cl2CH2], 77 [ClCHCOH], 48 [ClCH].

Dimethyl phosphate and dichloroacetic acid were novel metabolites only present in the DDVP degradation pathway. 2,2-dichloroethanol and phosphoric acid were metabolites existing in strain T23 growth process, but their content was significant increase in DDVP degradation pathway. The peak indicated dichloroacetic acid subsequently disappeared after ten days of incubation, indicating that these metabolites were finally degraded. Potential catabolic pathway for DDVP degradation by strain T23 was shown in Fig. 3C. The results of degradation experiments and metabolite identi?cation indicated that strain T23 could utilize DDVP through carbon co-metabolism in Burk medium and completely mineralized DDVP. Thus, in light of its broad-spectrum substrate speci?cities and complete mineralization of DDVP, strain T23 has the potential to be applied to the removal of DDVP and its metabolite residues from the environment.

**Fig. 3.**
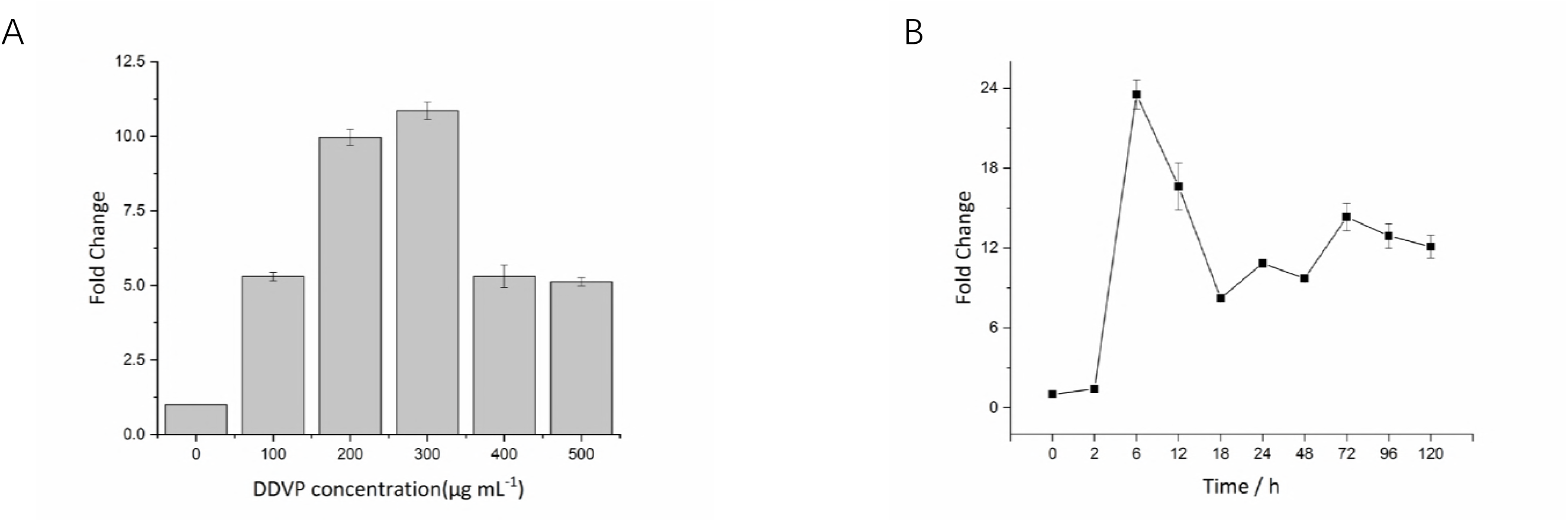
Expression analyses of *TaPon1*-like in *T. atroviride* T23. (A) *TaPon1*-like gene expression in liquid Burk medium with different concentrations of DDVP (100-500 μg mL^−1^) at 24 h. Medium without DDVP was used as a control. The graph shows the averages and means ± standard errors of three replicates. (B) Time course of *TaPon1*-like expression at 300 μg mL^−1^ DDVP. Time points on the *x-axis* represent the duration of DDVP stress. Relative mRNA levels were normalized according to *actin* gene expression and were calculated using the 2^−^ΔΔ^Ct^ method.

Gene expression analysis of *TaPon1*-like in *T. atroviride*

Gene expression analysis using a reverse transcription quantitative polymerase chain reaction (RT-qPCR) revealed a significant (*P* < 0.001) induction of *TaPon1*-like with different DDVP concentrations at 24 h (Fig. 3A). In addition, expression of *TaPon1*-like was strongly induced (*P* < 0.001) when T23 was exposed to 300 μg mL^−1^ DDVP. As shown in Fig. 3B, the expression of *TaPon1*-like significantly reached its maximum at 6 h and expressed a continuously high level during DDVP degradation process at an initial DDVP concentration of 300 μg mL^−1^ in liquid Burk medium.

Cloning and sequence analysis of *TaPon1-*like

*TaPon1-*like cDNA was amplified from the *T. atroviride* strain T23 genome. *TaPon1-*like was 1384 bp in length, including a 67-bp intron that encodes 438 amino acid residues. A neighbor-joining phylogenetic tree was constructed based on the amino acid sequence of TAPON1-LIKE and enzymes reported in GenBank, including hydrolases derived from bacteria and mammals (Fig. 4), to infer evolutionary relationships. A BLAST analysis showed that TAPON1-LIKE shared 10% to 21.83% identity to paraoxonase, arylesterases, and lactonases.

**Fig. 4.**
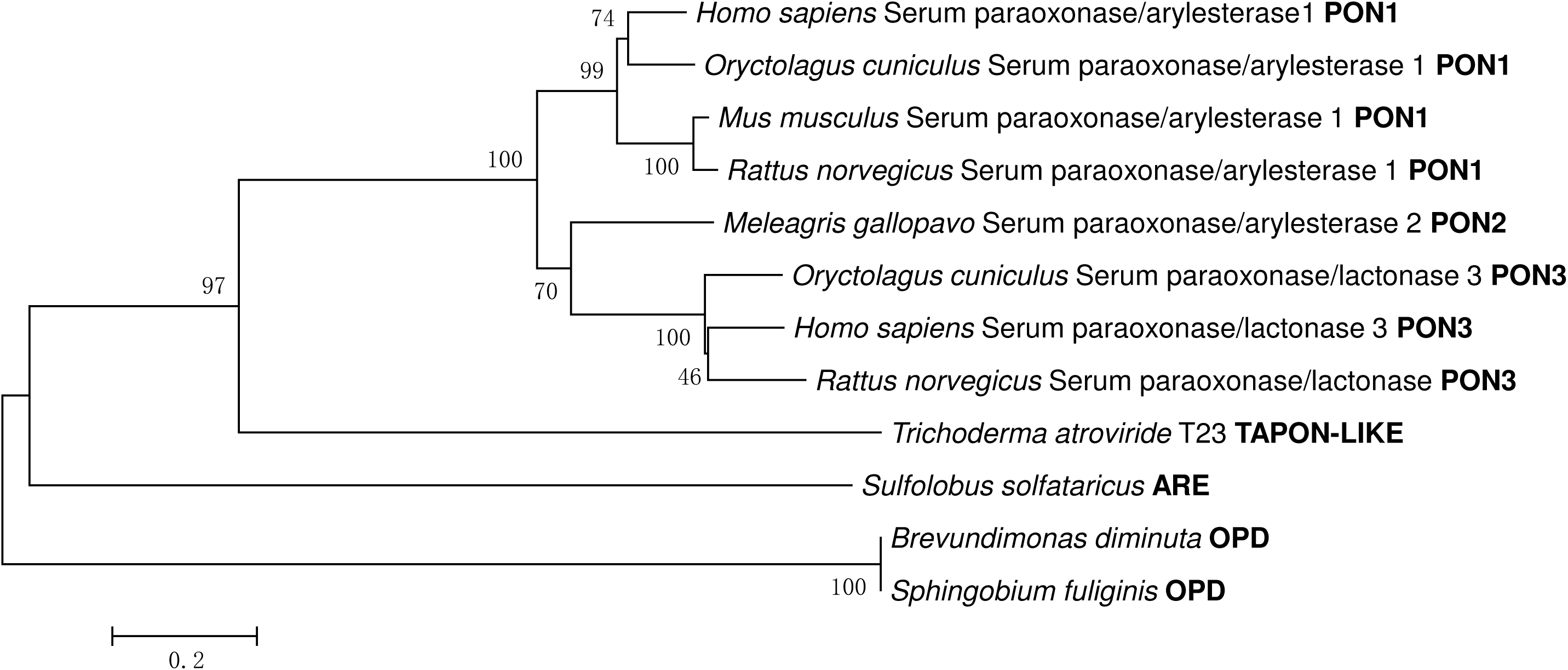
Phylogenetic tree of TAPON1-LIKE and related paraoxonases, arylesterases, and hydrolases constructed using the neighbor-joining method. The hydrolases and their accession numbers: HuPON1 (P27169), MosPON1 (P52430), RabPON1 (P27170), RatPON1 (P55159), BdOPD (P0A434), SsARE (B5BLW5), BdOPD (P0A434), SfOPD (P0A433), MosPON2 (Q91090), HuPON3 (Q15166), RabPON3 (Q9BGN0), and RatPON3 (Q68FP2).

Further analysis showed that TAPON1-LIKE shared 21.36% to 21.83% identity with HuPON1 proteins in the PDB database (PDB accession numbers 4Q1U.1.A, 3SRE.1.A, 4HHQ.1.A, AND 4HHO.1.A) such as the calcium-dependent lactonase that also promiscuously catalyzes the hydrolysis of paraoxon.

HuPON1 is a six-blade β-propeller containing two calcium ions in a central tunnel. The tunnel-buried calcium is critical for the enzyme’s conformational stability, and the solvent-exposed calcium residing at the bottom of the active-site cavity is needed for catalysis (14). In comparison with HuPON1, the mimetic TAPON1-LIKE contained Asn168, which ligates the catalytic calcium, while the Asp183-His184 dyad is vital for stabilizing the catalytic calcium ion.

Moreover, TAPON1-LIKE was found to have a significant motif, XXXTLVDNXXXXD, which may be the direct catalytic calcium metal-binding region for activating hydrolysis of OPPs, which interact with the side chains of Asp269 and Glu53. Motifs PXXPXXIXLMD or DXXXXXXXXMYLXVVN may be another metal-binding regions for supposed catalytic calcium ion (15) (Fig. 5 and Fig. S4C).

**Fig. 5.**
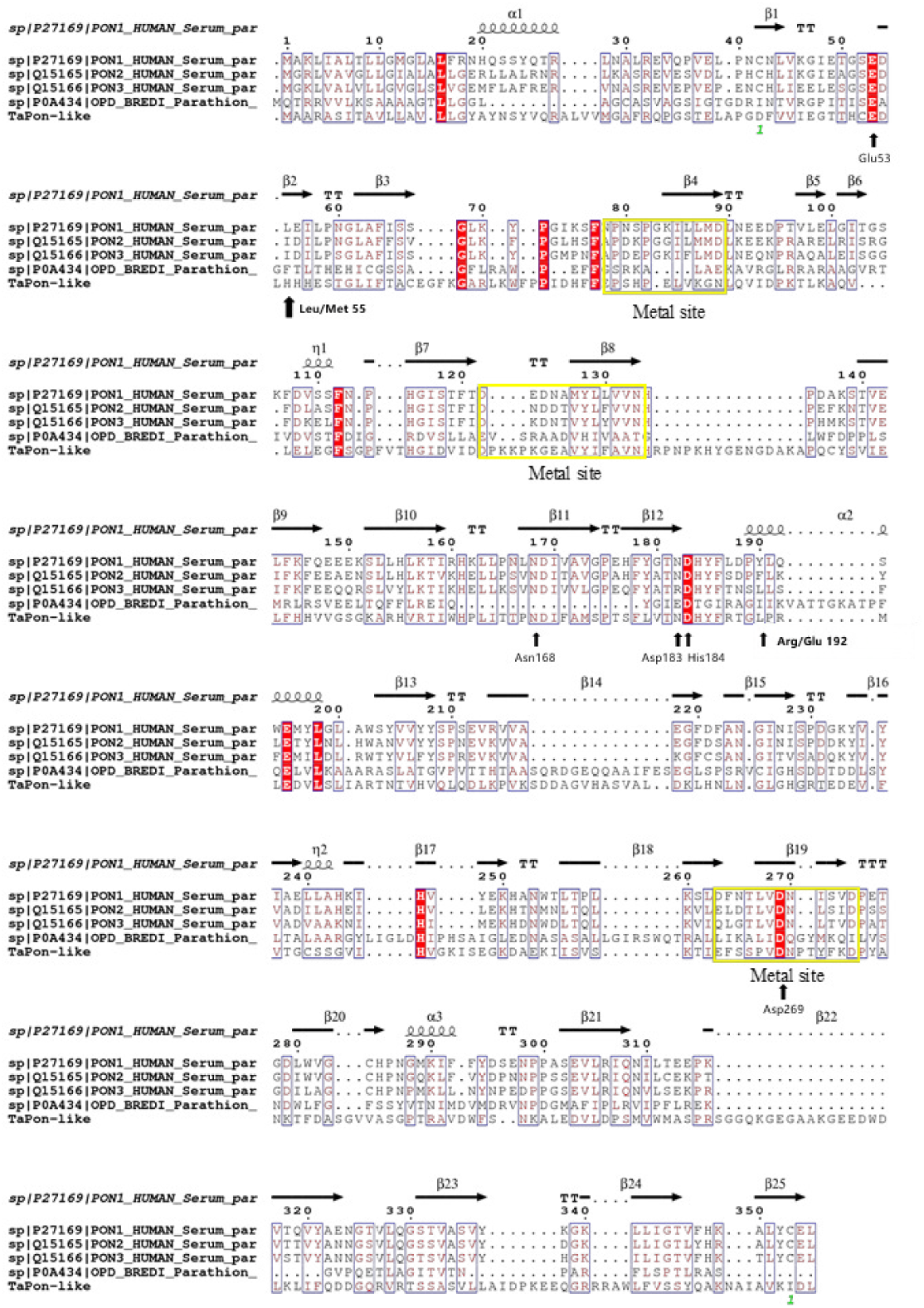
Multiple alignments of amino acid sequences of TAPON1-LIKE and similar proteins. The deduced amino acid sequences of human PON1, human PON2, human PON3, OPD (*Brevundimonas diminuta*), and TAPON1-LIKE are shown, respectively, from the first line to the bottom line. For the human species, there are two polymorphic sites that are indicated above at positions 55 and 192. Three yellow, boxed regions represent potential calcium-binding regions; the arrows represent conserved amino acid residues.

*TaPon1-*like function in DDVP degradation

To verify the function of the *TaPon1-*like gene in DDVP degradation, knockout mutants and complementation mutants were constructed. *TaPon1*-like knockout mutants were generated by replacing *TaPon1-*like with the *hygB* selection cassette by homologous recombination using ATMT (Fig. S2A). Twenty-seven hygromycin-resistant T23 colonies were obtained on selection plates containing timentin (300 μg mL^−1^) and hygromycin (200 μg mL^−1^). The individual transformants were subcultured on fresh selection plates, and eight hygromycin-resistant colonies were randomly selected for mutant validation to confirm that the knockout cassette was inserted at the correct position. *TaPon1-*like fragments were lost in top six deleted transformants but were observed in wild-type T23 (Fig. S2B). The flanking fragment from six transformants adjacent to the upper border demonstrated that the T-DNA insertion position of the knockout was correct (Fig. S2C). In addition, an RT-PCR experiment using primers specific to the *TaPon1*-like sequence demonstrated the complete loss of the *TaPon1*-like transcript in six transformants, while an amplification product of the desired size was found in the WT (Fig. S2D). Southern blotting (Fig. S2F) results showed *TaPon1*-like was replaced using an 800 bp fragment of *hyg*B. We selected one suitable deletion transformant designated KO1. Using similar methods, *TaPon1*-like complementation transformants were screened, and the suitable one was designated CO1 (Fig. S3).

The DDVP degradation rates of strain T23 and related mutants were measured by GC-FPD at an initial concentration of 300 μg mL^−1^ in Burk medium for 168 hours (h). Strain T23 exhibited an excellent efficiency of degrading DDVP, and the degradation rate value was 56% at the first 24 h (Fig. 6). CO1 presented a similar degradation rate curve compared with strain T23, and meanwhile no DDVP was residual in treatment by strain and CO1. KO1 declined in the DDVP biodegradation rate about 31% at 24 h compared with strain T23. The self-degradation rate of DDVP is almost 3% per 24 h. On the basis of between strain T23 and related mutants, *TaPon1-*like was shown to be a key gene in DDVP biodegradation.

**Fig. 6.**
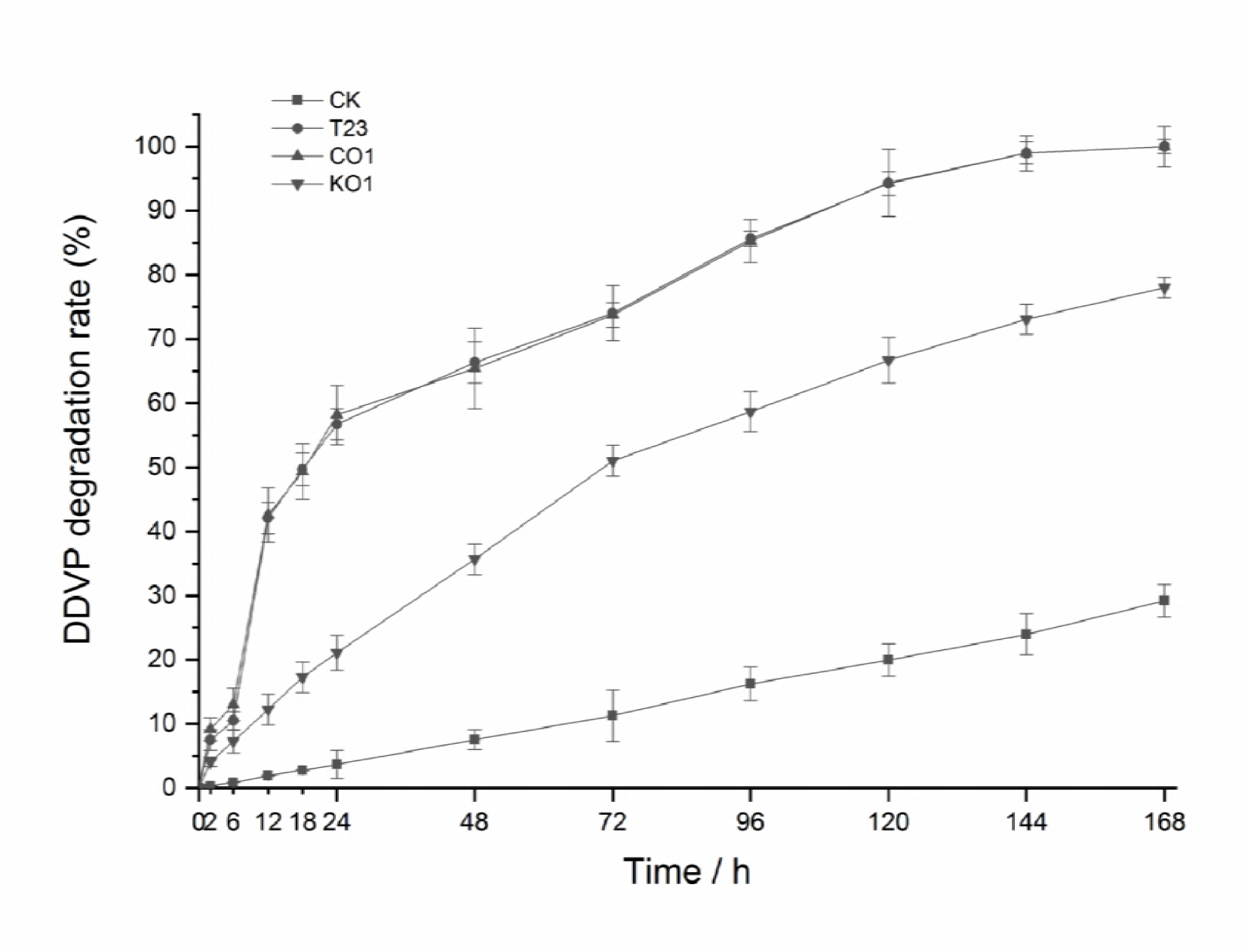
Degradation rate of DDVP by T23 and *TaPon1-like* mutants. Degradation rate curves of DDVP by mycelia of T23 and *TaPon1-like* mutants. The initial concentration of DDVP was 300 μg mL^−1^. CK, Burk medium with DDVP only; T23, wild type strain; CO1, *TaPon1-like* complementation mutant; KO1, *TaPon1-like* knockout mutant. Data are expressed as the means±standard errors of three replicates.

Sun et al. (16) showed that among 110 genetically stable T-DNA transformants of *T. atroviride* T23, one transformant, AMT-12, was confirmed by Southern blot analysis to have single-copy inserts of T-DNA, had 10% greater DDVP-degradation capacity than the wild type, and tolerated up to 800 μg mL^−1^ DDVP. Based on the changes in fungal biomass, gene expression and the variation in the biodegradation rate, we presumed that *TaPon1*-like played an important role in the DDVP degradation pathway.

Heterologous expression and purification of TAPON1-LIKE

Bioinformatics predicted that the isoelectric point of TAPON1-LIKE protein was 6.64 (Fig. S4A and Fig. S4B). TAPON1-LIKE had one secretion signal peptide at the N terminus, and there was no transmembrane domain in the protein (Fig. S4D). We determined whether the TAPON1-like protein contributed to DDVP degradation by determining the activity of a recombinant TAPON1-LIKE construct (with a 30-aa secretion signal peptide removed and the remaining fragment of the ORF cloned into pGEX-4T-1) produced by *E. coli* Origami B (DE3).

When TAPON1-LIKE was expressed in soluble form in *E. coli* Origami B (DE3) using one vector, the GST-tagged (26 kDa) TAPON1-LIKE protein was predicted to appear at approximately 71 kDa. According to the SDS-PAGE analyses, the purified enzyme produced a single band and was designated reTAPONN1-LIKE (Fig. 7).

**Fig. 7.**
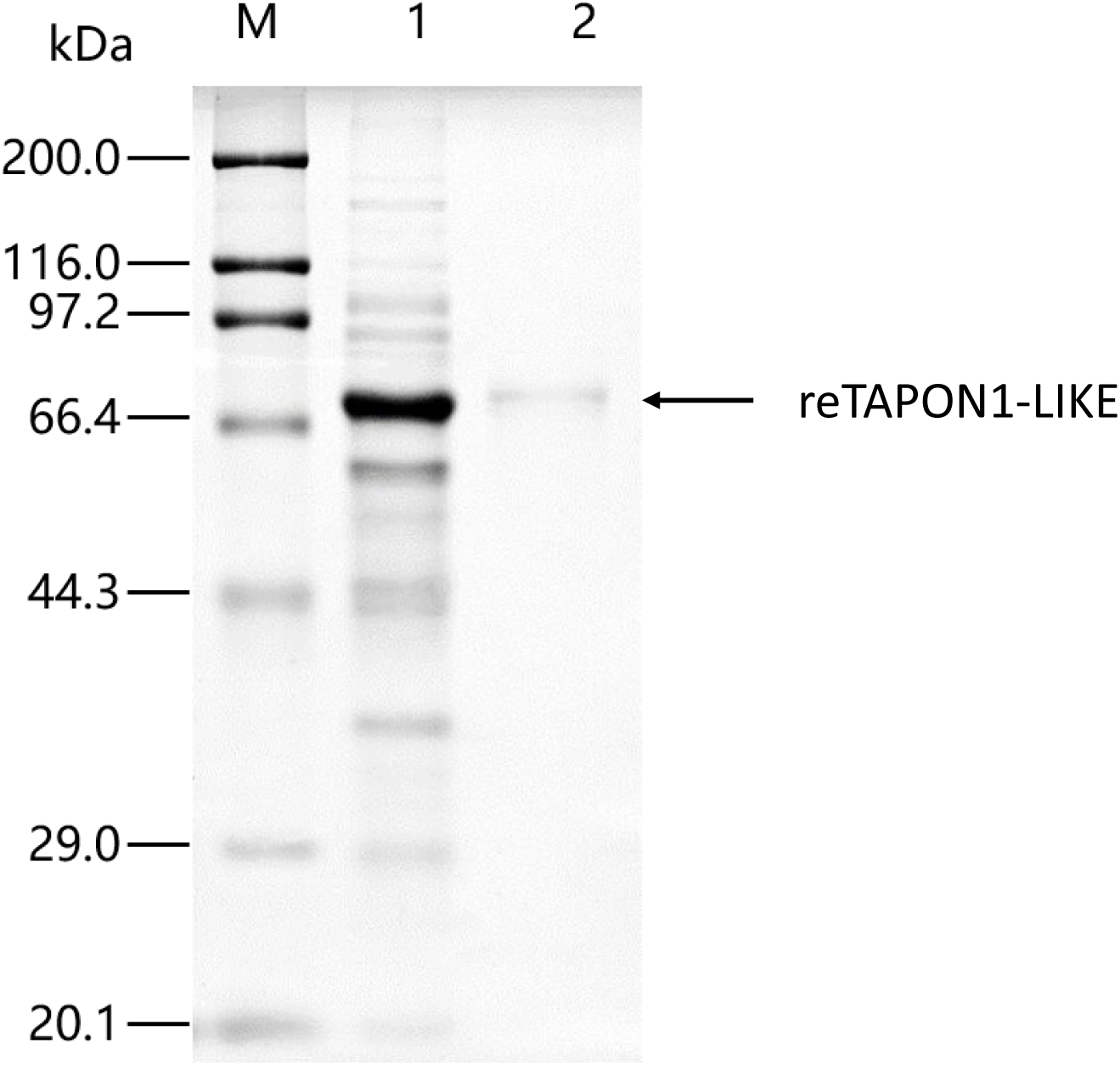
SDS-PAGE analysis of reTAPON1-LIKE heterologs expressed in the *E. coli* recombinant system.Lane M: protein marker, lane 1: supernatant protein of *E. coli* Origami B(DE3) harboring reTAPON1-LIKE induced by IPTG, lane 2: purification of reTAPON1-LIKE, with a predicted molecular mass of approximately 71 kDa.

The DDVP degradation products of purified reTAPON1-LIKE were evaluated by GC-MS analysis. Only *tert*-butyldimethylsilyl derivative of dimethyl phosphate, with a retention time of 14.106 min, was detected as a metabolite of DDVP biodegradation (Fig. S5). Therefore, we confirmed that TAPON1-LIKE is the enzyme to transform DDVP into dimethyl phosphate.

“PON1 activity” can be measured using substrates such as paraoxon, phenyl acetate, 4-nitrophenyl acetate, 5-thiobutylbutyrolactone (TBBL), and dihydrocoumarin (17), and the substrates of above could be classified into three types as paraoxon, aryl esters, and lactones. ReTAPON1-LIKE exhibited paraoxonase activity against DDVP (1028±31 U L^−1^) and chlorpyrifos-oxon (8198±53 U L^−1^). The arylesterase activity against p-nitrophenyl acetate and phenyl acetate was 6.16±0.02 U mL^−1^. The lactonase activity against hydrocoumarin was 10.89±0.06 U mL^−1^.

The purified protein or plasma derived from human and rabbit sera has been demonstrated to catalyze the hydrolysis of a broad range of substrates, including some pesticides such as oxon, arylesters of carboxylic acids, and lactones of hydroxy acids (18,20). Purified rabbit serum was injected into the tail veins of rats, increasing the peak hydrolytic activity of rat serum toward paraoxon by 9-fold and increasing that toward chlorpyrifos-oxon by 50-fold (21). The degradation of human fresh-frozen plasma was rapid, with half-lives of 19.5 s for chlorpyrifos-oxon and 17.9 min for DDVP (22). Engineered HuPON1 in the *E. coli* expression system showed kinetic parameters (*Km* and *V*max) of 0.121-0.317 mM and 34.7-245 U mg^−1^ protein for chlorpyrifos-oxon and 0.957-3.22 mM and 3020-680 U mg^−1^ for phenyl acetate (23, 24). Although the arylesterase and lactonase activities of reTAPON1-LIKE were decreased, we considered the superior catalytic efficiency against organophosphate-oxons pesticides. Comprehensive analysis confirmed that TAPON1-LIKE was considered PON-like, with superior catalytic efficiency against organophosphate-oxons, particularly DDVP.

Biochemical properties of reTAPON1-LIKE

The substrate specificities of reTAPON1-LIKE were determined by examining its activity against DDVP, chlorpyrifos-oxon, mevinphos, malaoxon, omethoate, methamidophos, chlorpyrifos, triazophos, phenyl acetate, p-nitrophenyl acetate, p-nitrophenyl butyrate, and 3,4-dihydrocoumarin (Table 1). Substrate spectrum analysis revealed that reTAPON1-LIKE nonspecifically reacted with substrates and that the enzyme could hydrolyze organophosphates, esters, and lactones. The most suitable substrate for reTAPON1-LIKE was P=O phosphotriester group pesticides, such as DDVP, chlorpyrifos-oxon, and mevinphos, with *Km* values of 0.23, 0.32, and 0.44 mM, respectively. Dimethyl phosphate pesticides (another group of organophosphates) were hydrolyzed by reTAPON1-LIKE but at low catalytic efficiency (≈10^3^ s^−1^ M^−1^). However, reTAPON1-LIKE revealed no activity against methamidophos, chlorpyrifos, and triazophos, which belong to the P=S phosphotriester group and methyl phosphate pesticides. The *Km* values for ester substrates were very similar to those of HuPON1 (25). The *kcat* values indicated that the catalytic efficiency of reTAPON1-LIKE against arylesters is determined by the carbon chain length of the acyl group. The reTAPON1-LIKE hydrolyzed 3,4-dihydrocoumarin, which is the most commonly used lactone, at a high catalytic efficiency (4.47×10^6^ s^−1^ M^−1^).

**Table 1.** Kinetic analysis of substrate hydrolysis by the reTAPON1-LIKE enzyme

| Substrate | $K_m$<br>(mM) | $k_{cat}$<br>$s^{-1}$ | Catalytic<br>efficiency<br>($k_{cat}/K_m s^{-1} M^{-1}$) |
| --- | --- | --- | --- |
| Dichlorvos | 0.23±0.02 | 204.3±2.0 | 8.88×10<sup>5</sup> |
| Chlorpyrifos-oxon | 0.32±0.05 | 1936.0±51.6 | 6.05×10<sup>6</sup> |
| Mevinphos | 0.44±0.04 | 6.5±0.3 | 1.47×10<sup>4</sup> |
| Malaoxon | 0.72±0.01 | 9.7±0.4 | 1.35×10<sup>4</sup> |
| Omethoate | 0.79±0.12 | 6.1±0.1 | 7.72×10<sup>3</sup> |
| Methamidophos | nd | nd | nd |
| Chlorpyrifos | nd | nd | nd |
| Triazophos | nd | nd | nd |
| Phenyl acetate | 0.89±0.06 | 4535.2±22.6 | 5.10×10<sup>6</sup> |
| p-Nitrophenyl acetate | 1.12±0.14 | 211.4±10.9 | 1.98×10<sup>5</sup> |
| p-Nitrophenyl butyrate | 1.58±0.12 | 59.8±1.1 | 3.78×10<sup>4</sup> |
| 3,4-Dihydrocoumarin | 0.73±0.01 | 3261.1±73.0 | 4.47×10<sup>6</sup> |

Note: Each value corresponds to the average of three different determinations. nd, not determined.

Enzyme dynamics of reTAPON1-LIKE

A variety of environmental factors such as pH and temperature exert varied effects on the enzyme activity and stability (6). The reTAPON1-LIKE protein exhibited high activity to DDVP between pH 6.5 and 10.0 (more than 50% relative activity), and pH 8.5 was optimum for its activity (Fig. 8A). Serum paraoxonase 1 activity was reported to occur from pH 7 to 10.5 (25), and the ideal pH of the reaction is in the range of 8 to 8.5 (26, 27).

**Fig. 8.**
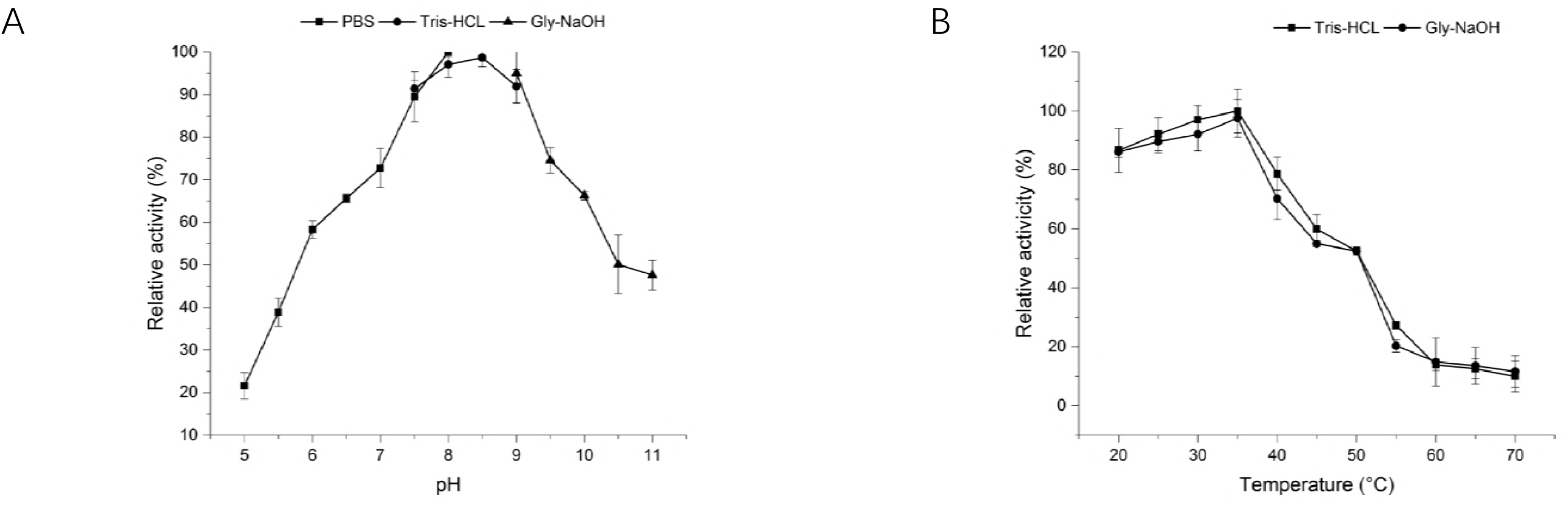
Effects of pH (A) and temperature (B) on reTAPON1-LIKE enzymatic activity. DDVP was used as the substrate. Activity at the optimal pH and temperature was defined as 100%. Data are expressed as the means ± standard errors of three replicates.

The reTAPON1-LIKE protein was active at 20-50 °C, with an optimum temperature of 35 °C using DDVP as substrate (Fig. 8B). An increase in temperature of 1 °C was associated with a 4.5% increase in PON1 activity when phenyl acetate was used as substrate (28). The optimum temperature of hydrolysis for paraoxon and phenyl acetate ranges from 30-45 °C (29).

The TAPON1-LIKE amino acid sequence shared identity with the sequences of the metal-dependent hydrolases; therefore, it was deduced that metal ions might also affect reTAPON1-LIKE enzymatic activity. As shown in Table 2, ethylenediaminetetraacetic acid (EDTA) inhibited the enzymatic activity at a final concentration of 1%; the result indicated that the reTAPON1-LIKE activity might be that of a metal ion-dependent hydrolase.

**Table 2.** Effects of different metal ions on the activity of the reTAPON1-LIKE enzyme

| Additive | Concentration (mM) | Relative activity (%) |
| --- | --- | --- |
| No addition |  | 100±1.5 |
| Metal ions |  |  |
| Ca<sup>2+</sup> | 1 | 589.7±3.3 |
| Zn<sup>2+</sup> | 1 | 234.4±5.6 |
| Mg<sup>2+</sup> | 1 | 121.2±4.6 |
| Cu<sup>2+</sup> | 1 | <5 |
| Ni<sup>2+</sup> | 1 | 95.5±2.4 |
| Na<sup>+</sup> | 1 | 176.5±5.2 |
| Ba<sup>2+</sup> | 1 | 151.6±1.4 |
| Mn<sup>2+</sup> | 1 | 119.9±0.8 |
| Fe<sup>2+</sup> | 1 | 93.2±0.6 |
| Co<sup>2+</sup> | 1 | 82.7±1.8 |
| Enzyme inhibitors |  |  |
| EDTA | 1 | nd |

Note: The activities of metal ion-treated reTAPON1-LIKE enzyme were assayed under standard assay conditions after incubation with 1 mM of various metal ions. The activity of the reTAPON1-LIKE enzyme with no added metal ions was set as 100%. Each value corresponds to the average of three different determinations.

As can be seen in Table 2, incubation of Ca^2+^, Zn^2+^, Na^+^ and Ba^2+^ resulted in complete reactivation to 589.7% ± 3.3%, 234.4% ± 5.6%, 176.5% ± 5.2%, and 151.6% ± 1.4%, respectively. On the contrary, the Cu^2+^ and Co^2+^ ions had inhibited the TAPON-LIKE activity toward DDVP at a final concentration of 1.0 mM L^−1^. PON1, whether it is derived from rabbit serum or human serum, its catalytic activity was dependent on Ca^2+^ ion. In addition, the stability of human PON1 was also dependent on Ca^2+^ ion, which corresponds to the two Ca^2+^ active centers in the three-dimensional structure of PON1 (3). Some ions (such as Zn^2+^, Co^2+^, Mn^2+^, Sr^2+^, Ba^2+^, Cd^2+^ or Mg^2+^) were noted as the protector to keep the human PON in an active form (30). However, the purified reTAPON1-LIKE was more like phosphodiesterase isolated from bacteria *Sphingobium* sp. and *Flavobacterium* sp., which could recovery the activity by obtaining these metal ions (Zn^2+^, Mn^2+^, Ba^2+^,Cd^2+^ or Mg^2+^). Moreover, (31) reported the hydrolysis of tetriso by an enzyme derived from *Pseudomonas diminuta* as a model for the detoxication of O-ethyl S-(2-diisopropylaminoethyl) methylphosphono; and (32) demonstrated the structure of a novel phosphotriesterase from *Sphingobium* sp. TCM1, which had a familiar binuclear metal center embedded in a seven-bladed β-propeller protein fold. Thus, calcium ion plays an important role in the catalytic activity of reTAPON1-LIKE3 to DDVP, but Ca^2+^, Zn^2+^, Na^+^ and Ba^2+^ also serves to stabilize the enzyme by keeping its native molecular structure and reactivate the catalytic activity.

## Discussion

The secrets of how DDVP is degraded by *T. atroviride* T23 and the intra- and extracellular enzymes involved in the DDVP biodegradation are still big challenges. Even though a group of *Trichoderma* spp. strains has been widely applied in bioremediation of chemical pesticide-polluted environments, until now, only a few genes, such *hex1* (10) and *TaPdr2* (11), have been found to play roles in the microbial tolerance to stress.

PON1s derived from mammals such humans, rabbits, and rats demonstrated the metabolic function of removing organophosphate pesticide residues from blood (33). Degradation of human fresh-frozen plasma containing high levels of HuPON1 was rapid, with the shortest half-life of 17.9 min for DDVP (22). Injection of PON1 into rats with acute organophosphate poisoning can decrease the amount of DDVP that enters the blood, lower the peak concentration, and relieve clinical signs(34).

Since previous researches have already revealed that serum PON1 is responsible for the degradation of DDVP, we hypothesized that a similar mechanism of DDVP biodegradation also present in *Trichoderma*. In our study, an effective protein, designated TAPON1-LIKE, with biodegradation activity of DDVP, was demonstrated to be a hydrolase. The gene encoding TAPON1-LIKE included a 1317-bp ORF, and the deduced amino acid sequence shared a certain homology with HuPON1, which may also have two calcium-binding sites. Using the ATMT method, mutants were constructed, and the function of the *TaPon1-like* gene in the degradation of DDVP was verified. Expression and purification of recombinant enzyme is a way to understand the properties of TAPON1-LIKE. reTAPON1-LIKE showed broad activity ranges for substrate, temperature, and pH. In addition, stimulating and inhibiting metal ions, optimum electron donors, and kinetic parameters were identified.

Human cDNA clones revealed that PON1 has two common coding polymorphisms, L55M and Q192R. Some studies have shown that genetic polymorphisms of the PON1 192 site can influence the activity of PON1, which may modify the individual susceptibility to methylparathion-induced toxicity (effects of PON1 polymorphism on the activity of serous PON in workers exposed to organophosphorus pesticides) and the catalytic efficiency of hydrolysis of paraoxon and chlorpyrifos oxon(35). In our study, the sequencing results for *TaPon1*-like verified that the polymorphism of the TAPON1-LIKE 192 site involved residue Arg192, similar to human PON1, which may determine the high catalytic efficiency.

The recombinant enzyme was also confirmed to have some of PON1 activity in the biodegradation of DDVP. Similarly, the TAPON1-LIKE expressed in *E. coli* showed different activities on a range of substrates, and this result suggested that the *TaPon1*-like gene was involved in the biodegradation of different paraoxon-like pesticides, especially for significantly improving the efficiency of pesticide-oxons such as chlorpyrifos-oxon and mevinphos. It has been clearly shown that paraoxon, chlorpyrifos-oxon, and DDVP can all be hydrolyzed by purified HuPON1; in addition, HuPON1 is able to function on a ranges of substrates, such as phenyl acetate, 4-nitrophenyl acetate, TBBL or dihydrocoumarin, chlorpyrifos, diazinon, sarin, or soman, among others(36). It was further found that reTAPON1-like with a broad substrate spectrum towards three types of catalytic substrate, but the enzyme activities showed some differences in kinetic parameters compared with HuPON1. For example, the reTAPON1-LIKE catalytic efficiency for phenyl acetate and 3,4-Dihydrocoumarinpesticide was 5.10×10^6^ and 4.47×10^6^, which were lower than HuPON1 (26).

In conclusion, this study found that *T. atroviride* strain T23 produced TAPON1-LIKE protein with functions in the biodegradation of DDVP. The more valuable of this work were provided a novel clue to comprehensively understanding the degradation mechanism of a series of residual organophosphate pesticides through Trichoderma which has been widely applied as bioremediation approach worldwide. The protein’s differential roles in the biodegradation of various organophosphate pesticides and its expression and properties in other Trichoderma species remain under study.

## Materials and Methods

Reagents and media

Main chemical reagents such as pesticide standards, including DDVP, chlorpyrifos, mevinphos, malaoxon, omethoate, triazophos, and methamidophos, were purchased from Dr. Ehrenstorfer GmbH (Augsburg, Germany). Chlorpyrifos-oxon, p-nitrophenyl acetate, p-nitrophenyl butyrate, phenyl acetate, and 3,4-dihydrocoumarin were purchased from J&K Scientific Ltd. (Beijing, China). Other chemical reagents were purchased from the Sinopharm Chemical Reagent Co. Ltd. (Beijing, China). Difco PDA medium was purchased from Becton Dickinson & Co. (Franklin Lakes, NJ, USA). *N-tert*-Butyldimethylsilyl-*N*-methyltrifluoroacetamide (MTBSTFA) was purchased from Sigma (St. Louis, MO, USA). Burk medium contained (liter^−1^) 0.2 g K2HPO4, 0.8 g KH2PO4, 0.2 g MgSO4•7H2O, 0.1 g CaSO4•2H2O, 0.0033 g Na2MoO4•2H2O, 0.005 g FeSO4•7H2O, 1 g (NH4)2SO4, and 1 g glucose, pH 6.0. Luria-Bertani (LB), YEP, CYA, and IM media were prepared as previously described (37).

Strains, plasmids, and culture conditions

The strains, plasmids, and primers used in this study are listed in Table 3 and Table 4. The wild-type strain T23 and transformants were maintained in PDA (Difco, Becton Dickinson & Co., USA) at 28 °C until sporulation occurred. *Agrobacterium tumefaciens* strains were grown on YEP agar or in YEB broth at 28°C. *Escherichia coli* strains were grown in LB broth or LB agar plates. A knockout plasmid, pC1300qh, containing the hygromycin (hygB) resistance gene ORF, was constructed with a pCAMBIA1300 plasmid backbone in which the 35S promoter was replaced with the trpC promoter (37).

**Table 3.** Strains and plasmids used in this study

| Strain or plasmid | Description <sup>a</sup> | Reference or source |
| --- | --- | --- |
| Strains |  |  |
| *T. atroviride* T23 | Wild-type DDVP degrader | Laboratory stock |
| *T. atroviride* KO1 | Derivative of strain T23, site-directed knockout mutant of *TaPon1*-like ORF cassette, Hyg<sup>r</sup> | This study |
| *T. atroviride* CO1 | Derivative of KO 5, complementation transformant of *TaPon1*-like ORF cassette, G418<sup>r</sup> | This study |
| *A. tumefaciens* AGL1 | Host strain for *Agrobacterium tumefaciens*-mediated transformation of filamentous fungi, Rif<sup>r</sup> | Laboratory stock |
| *E. coli* DH5α | Host strain for cloning vectors | TaKaRa |
| *E. coli* Origami B (DE3) | Host strain for expression vector | Novagen |
| Plasmids |  |  |
| pGEX-4T-1 | Expression vector; Amp<sup>r</sup> | Novagen |
| pGEX-4T-1-*TaPon11*-like | pGEX-4T-1 harboring the *TaPon1*-like gene with signal peptide deleted | This study |
| pMD19-T | T-A cloning vector, Amp<sup>r</sup> | TaKaRa |
| pSilent-1 | Vector including trpC promoter and terminator sequences, Amp<sup>r</sup> | Laboratory stock |
| pC1300QH | Knockout broad-host-range shuttle vector, Kan<sup>r</sup> | (46) |
| pC1300QH-*TaPon11*-like | pC1300QH harboring the 5' flanking fragment and 3' flanking fragment of the gene *TaPon*-like, Kan<sup>r</sup> | This study |
| pC1300TH | Complementation broad-host-range shuttle vector, Kan<sup>r</sup> | (46) |
| pC1300TH-*TaPon1*-like | pC1300TH harboring the *TaPon*-like ORF complementation cassette, Kan<sup>r</sup> | This study |

<sup>a</sup> Amp<sup>r</sup>, ampicillin resistant; Kan<sup>r</sup>, kanamycin resistant; Rif, rifampin resistant; Hyg<sup>r</sup>, hygromycin B; G418<sup>r</sup>, geneticin resistant.

**Table 4.**
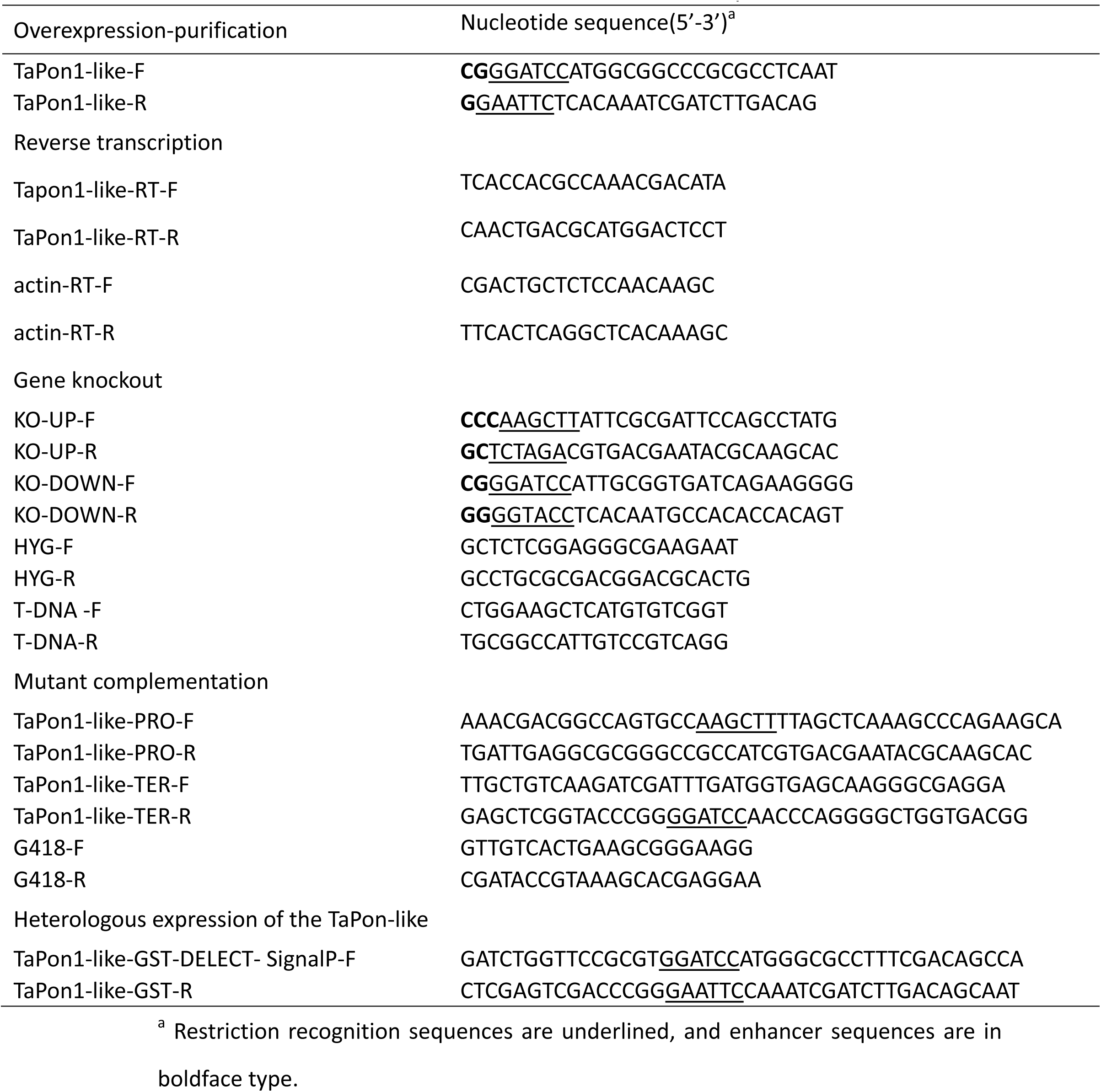
Primers used in this study

Growth and degradation experiments with fungal strains

In earlier studies it was observed that fungus played a significant role in reducing organic compound levels through enzymes they produce or mycelial adsorption (38), (39), (40), (41). Thus, it was decided to study the degradation and adsorption performed by strain T23. Agar plugs of strain T23 and *TaPon1-*like mutants were precultivated for 2 days in 300 mL flasks containing 500 mL of PD medium on a rotary shaker (180 rpm) at 28 °C, followed by harvesting of mycelia by filtering these cultures through filter paper. The harvested mycelia were washed three times with sterile distilled water, and then 1 g of wet mycelia was transferred to new flasks containing 50 mL of Burk medium. Mycelia transferred in fresh Burk medium without DDVP was used as a control.

We examined the degradation and adsorption of two DDVPs by strain T23 under two sets of conditions: (a) at ten time points (0-120 h) with an initial concentration of 300 μg mL^−1^ DDVP and (b) different initial concentrations of DDVP (100-500 μg mL^−1^) at 24 h.

The biomass of strain T23 was determined by measuring by the dry weight of mycelia after vacuum drying at −40 °C for 12 h in an Alpha freeze-dryer (Christ, Osterode, Germany). The DDVP concentration in Burk medium and adsorbate concentration of DDVP were assessed using a GC-2010-FPD Plus (Shimadzu, Japan) according to the methods described by Xiao et al. (42). To quantify the absorption of DDVP by mycelia, the dried mycelia were treated with spray gold and then analyzed by scanning electron microscopy (SEM, NOVA NanoSEM 230, FEI, USA) equipped with energy dispersive spectroscopy (EDS, Aztec X-Max, Oxford Instruments, UK). All treatments consisted of three replicates.

Homologous cloning of *TaPon1-*like genes in *T. atroviride* T23

We queried the *T. atroviride* genome v2.0 (IMI 206040) on JGI (https://genome.jgi.doe.gov/Triat2/Triat2.home.html) and used local BLAST to search for genes based on the homology domain of *HuPon1* in the human liver. Subsequently, a homologous gene named *TaPon1-*like (GenBank accession number MH802589) was identified in *T. atroviride*. Strain T23 genomic DNA was used as a template for *TaPon1-*like gene amplification with the primers TaPon1-like-F and TaPon1-like-R. Total RNA of T23 was extracted from frozen powdered mycelia using TRIzol (11), and total cDNA of T23 was synthesized using the PrimeScript RT Reagent kit (TaKaRa, Dalian, China). The *TaPon1-*like ORF was PCR-amplified using the primers TaPon1-like-F and TaPon1-like-R. The PCR product was purified from a gel extraction, ligated into a PMD19-T vector, and then used as a PCR template in the assays described below. The recombinant plasmid was transformed into *E. coli* DH5α competent cells, and the cells were then isolated from the transformants and sequenced.

Comparison of the deduced amino acid sequences in the genes

The deduced amino acid sequences of *TaPon1-*like genes were compared with the amino acid sequences of similar genes in other species using PDB database search sequences (http://www.rcsb.org/#Subcategory-search_sequences). Amino acid sequence alignment was conducted using ClustalX (43) and Espript 3.0 (http://espript.ibcp.fr/ESPript/ESPript/). The phylogenetic tree was constructed using the neighbor-joining method and the software MEGA 5.0. A bootstrap consensus tree was inferred from 1000 replicates and represents the evolutionary history of the taxa analyzed. Signal peptides were analyzed using the SignalP 4.1 Server (http://www.cbs.dtu.dk/services/SignalP/), and the membrane-spanning domains were calculated using TMHMM v. 2.0 (http://www.cbs.dtu.dk/services/TMHMM/).

Gene expression analysis

Gene expression analysis of *TaPon1*-like genes in strain T23 was performed under the two different conditions described above. RNA extraction was performed using the Qiagen RNeasy kit following the manufacturer’s protocol (Qiagen, Hilden, Germany). One microgram of total RNA was reverse transcribed in a total volume of 20 μL as described above. Transcript levels were quantified by RT-qPCR using SYBR Green PCR Master Mix (TaKaRa, Dalian, China) and the primer pair TaPon-like-RT-F/TaPon-like-RT-R (Table 2) in a LightCycler^®^ 96 system (Roche, Basel, Switzerland). Relative expression levels for the target gene in relation to those of the *actin* gene using the primers actin-RT-F and actin-RT-R were calculated from the Cq values and the primer amplification efficiencies using a formula described previously (44). Gene expression analysis was performed in three biological replicates, with each based on three technical replicates.

Identification of DDVP degradation metabolites

Strain T23 (500 μg mL^−1^) was inoculated into a 500 mL Erlenmeyer flask containing 200 mL of Burk medium, and the culture was cultivated as described above. The metabolites were analyzed by gas chromatography-mass spectrometry (GC-MS). After an incubation period, the mycelia were removed by filtering through filter paper. Then, 2% NaCl and 2 mL of HCl were added; 100 mL of anhydrous ethyl ether was subsequently added (equal to 50% of the total volume), and the sample was mixed for 5 min on a vortex mixer. After the liquid stood for 10 min, the ether layer was collected thrice. The ether fractions were concentrated at 30 °C with a rotary evaporator and dried over anhydrous Na2SO4.Remaining aqueous phase was concentrated 100-fold at 30 °C with a rotary evaporator until no liquid was present. One of the moist precipitates was then extracted with two 3 mL portions of acetone. The combined acetone extract was concentrated by a stream of N2, and 100 μL derivative reagent MTBSTFA was added, and the solutions were mixed and heated for 60 min at 70 °C and were then ready for instrumental analysis

Qualitative and quantitative analyses were performed on a 7890A gas chromatograph and a 5975C mass spectrometer (Agilent Technologies, Milano, Italy). GC-MS instrumental conditions were similar to those of the assay of organophosphate compounds (OPPs) with the following operating parameters: HP-5 GC column (30 m×0.32 mm i.d., 0.50 μm film thickness); temperature program of a 60 °C initial temperature (6 min hold), 10 °C min^−1^ ramp to 250 °C, and 15 °C min^−1^ ramp to 280 °C (6 min hold). One microliter of each sample was injected into the GC-MS and analyzed using full scan mode, and the instrument was scanned from 10 to 600 amu.

Construction of knockout and complementation *TaPon1-*like mutants

To generate *TaPon1-*like knockout transformants, a homologous recombination cassette designated pC1300QH-TaPon1-like (Fig. S2A) was constructed with 928-bp 5’ flanking sequences and 896-bp 3’ flanking sequences using the primers KO-UP-F/KO-UP-R and KO-DOWN-F/KO-DOWN-R. The cloned flanking sequences were restriction-digested with *Hind*III, *Xba*I, *BamH*I, and *Kpn*I (Thermo Fisher Scientific, Shanghai, China), respectively, and these fragments were gel-purified and ligated to modified plasmid pC1300QH. *Agrobacterium tumefaciens*-mediated transformation (ATMT) was performed as previously described (37, 45) for the generation of *TaPon1-*like knockout and complementation transformants.

The genomic DNA of T23 and transformants was isolated using a modified cetyltrimethylammonium bromide (CTAB) method (46). To identify the *TaPon1-*like knockout transformants, (a) the *TaPon1-*like knockout was verified by attempting to amplify the *TaPon1-*like gene with the primers TaPon1-like-F and TaPon1-like-R; (b) the T-DNA insertion numbers were determined using the primers T-DNA–F and T-DNA–R; (c) RT-PCR analysis of *TaPon1*-like gene expression using Tapon1-like-RT-F/ Tapon1-like-RT-R primers. Finally, Southern blotting (47) using *hygB* as a probe was performed to confirm the knockout transformant.

The plasmid to complement the *TaPon1-*like knockout was constructed with the promoter, which was amplified using the T23 genome as a template with the primers TaPon1-like-PRO-F and TaPon1-like-PRO-R, the amplified *TaPon1-*like ORF, and the trpC terminator amplified from the vector pSilent-1 (48). We ligated the three PCR-amplified fragments into plasmid pC1300TH digested with *Hind*III and *BamH*I using a Hieff Clone™ Plus One Step Cloning Kit (Yeasen, Shanghai, China) to construct the *TaPon1-*like ORF complementary cassette, designated pC1300TH-TaPon1-like (Fig. S3), and then the ORF complementary cassette inserted into the genome of knockout mutant KO1.

Similarly, the complementary transformants were identified (a) the fragment of *TaPon1*-like gene fusion of *trp C* terminor using Tapon1-like-F/ Tapon1-like-TER-R primers, (b) the fragment of G418 gene amplified from genomic DNA using G418-F/ G418-R primers, (c) amplification of TAPON-LIKE complementation cassette from genomic DNA using Tapon1-like-PRO-F/ Tapon1-like-TER-R primers, and (d) RT-PCR analysis of *TaPon1*-like gene expression using Tapon1-like-RT-F/ Tapon1-like-RT-R primers.

Protein expression and purification of reTAPON1-LIKE

The expression plasmid pGEX-4T-1-*TaPon1-*like was constructed via the ligation of a partial gene sequence of the *TaPon1-*like gene that was lacking the 90-bp signal peptide into the corresponding restriction sites of a pGEX-4T-1T plasmid digested by *BamH*I/ *EcoR*I. The ligation was performed according to the Hieff Clone™ Plus One Step Cloning Kit manual. The expression plasmid pGEX-4T-1-*TaPon1-*like was then transformed into *E. coli* Origami B (DE3), and the recombinant purified protein was designated reTAPON1-LIKE.

The transformant was grown at 37 °C and 200 rpm in 1 mL of LB medium containing ampicillin (50 μg mL^−1^) until the OD600 reached 0.6, and then the culture was cultivated in 1 L of LB liquid medium and induced with 0.6 mM isopropyl β-D-1-thiogalactopyranoside (IPTG). After screening, the best DDVP degradation by the reTAPON1-LIKE expression strain was determined. The cells were harvested at 9000 rpm and 4 °C for 10 min and were then washed with PBS (phosphate buffer solution, 10 mM, pH 8.0). The pellets were suspended in lysis buffer: 0.2 mM phenylmethylsulfonyl fluoride (PMSF), 1 mM DL-dithiothreitol (DTT), 1 mM lysozyme, and 50 mM Tris-HCl. The mixture was sonicated with an ultrasonic cell disruptor (Jingxin Industrial Development Co., LTD, JY92-IIN, Shanghai, China) at 25 °C with 4- to 6-s cycle pulses for 30 min. The lysate containing the reTAPON1-LIKE fusion protein was centrifuged at 12,000 rpm and 4 °C for 20 min, and the supernatant was filtered through a 0.22-μm filter. The filtrate was loaded onto GST•Bind™ Resin (Novagen, Germany) pre-equilibrated with PBS. A flow rate of approximately 10 column volumes per hour was used. Then, the column was washed with 10 volumes of PBS, and the recombinant protein was eluted with three volumes of PBS containing 10 mM reduced glutathione at 4 °C. The target protein was concentrated, and reduced glutathione was removed. ReTAPON1-LIKE, including cell pellets and purified protein, was boiled for 5 min for denaturation, and its concentration was then estimated by SDS-PAGE.

Degradation of DDVP by purified reTAPON1-LIKE enzyme

The enzyme activity of purified reTAPON1-LIKE toward DDVP was measured for 30 min with 100 μM DDVP in a 50 mM glycine buffer, pH 8.5, containing 2.0 M NaCl and 1.0 mM CaCl2. DDVP and metabolites generated by reTAPON1-LIKE degradation activity in the reaction system were extracted with a 1/2 volume of anhydrous ethyl ether and mixed for 5 min at room temperature. The sample was centrifuged at 12,000 rpm, and the upper supernatant was collected in a 1.5 mL tube. This step was repeated twice, and the samples were then concentrated for GC-MS analysis as described above.

Substrate specificity of reTAPON1-LIKE and enzyme kinetics

The activity of HuPON1 can normally be measured using three types of substrates: paraoxon, unphosphorylated aryl esters, and lactones (17)

The paraoxonase activities of reTAPON1-LIKE were measured with paraoxonase-like pesticides such as DDVP (100-500 μM), chlorpyrifos-oxon (100-500 μM), mevinphos (10-100 μM), malaoxon (10-100 μM), omethoate (100-500 μM), triazophos (100-500 μM), and methamidophos (100-500 μM). The reaction conditions included a 50 mM glycine buffer, pH 8.5, containing 2.0 M NaCl and 1.0 mM CaCl2, as previously reported (49). The extraction of these compounds was performed according to the methods described above and detected by GC-FPD.

The arylesterase activity of reTAPON1-LIKE was determined with phenyl acetate (1.0-5.0 mM), p-nitrophenyl acetate (0.5-2.5 mM), and p-nitrophenyl butyrate (0.25-1.25 mM) as substrates in 20 mM Tris-HCl buffer, pH 8.0, containing 1.0 mM CaCl2, as previously described (50).

The lactonase activity of reTAPON1-LIKEs was determined with dihydrocoumarin (0.1-2.0 mM) as substrate in 25 mM Tris-HCl buffer, pH 7.4, containing 1.0 mM CaCl2, as previously described (50).

The arylesterase and lactonase activities of reTAPON1-LIKE were detected using a SpectraMax i3x Multi-Mode Detection Platform (Molecular Devices, CA, USA). Enzyme activities are expressed in international units (U) per liter (L) of supernatant enzyme reTAPON1-LIKE, and one unit is defined as the amount of enzyme that catalyzes the turnover of 1 μM of substrate per min at 25 °C.

The catalytic constants *k*_cat_ and *K*_m_ were determined under standard assay conditions with 7 different concentrations of various substrates and a range of reTAPON1-LIKE concentrations (0.05-5.00 μM). Kinetic parameters were determined using a Lineweaver-Burk plot.

Biochemical properties of purified reTAPON1-LIKE

The most desirable pH was determined by incubating the purified enzyme in phosphate buffer (pH 5.0-8.0), glycine-NaOH (pH 9.0-11.0), and Tris-HCl buffer (pH 7.5-9.0). To study the effect of temperature on the activity of purified reTAPON1-LIKE against DDVP at pH 8.0, the temperature of the assays was varied from 25 °C to 70 °C. Enzyme activity was reported as a percentage of the highest value, which was set as 100%.

The effects of metal ions and inhibitors on enzyme activity were investigated at 25 °C and pH 8.0. The purified reTAPON1-LIKE was incubated with 1 mM solutions of the metal salts MgCl2, MnCl2, ZnCl2, CaCl2, CuCl2, BaCl2, FeCl3, and NaCl for 10 min to determine residual activity. The results were expressed as percentages, and the values of the native enzyme without metal ion addition were set as 100%.

## Acknowledgments

This study was supported by the following projects: The National Key Research and Development Program of China (2017YFD0200403, 2017YFD0201108), the Natural Science Foundation of China (31672072; 31750110455), the Intergovernmental Key International Scientific and Technological Innovation Cooperation (2017YFE0104900), the 948 Project of the Ministry of Agriculture (2016-X48), and CARS-02. We all appreciate Lurong Xu for guiding us in the operation of the GC-MS instrument and analyzing the data.

## References

1. Gan Q, Singh RM, Wu T, Jans U. 2006. Kinetics and mechanism of degradation of dichlorvos in aqueous solutions containing reduced sulfur species. Environmental Science & Technology 40:5717–5723.

2. Liu C, Qiang Z, Adams C, Tian F, Zhang T. 2009. Kinetics and mechanism for degradation of dichlorvos by permanganate in drinking water treatment. Water Res 43:3435–42.

3. Lynch SM, Lorenz J, Klotz S. 2014. Inclusion of calcium during isolation of high-density lipoprotein from plasma maintains antioxidant function. Analytical Biochemistry 454:41–43.

4. Ren Z, Zhang X, Wang X, Qi P, Zhang B, Zeng Y, Fu R, Miao M. 2015. AChE inhibition: One dominant factor for swimming behavior changes of Daphnia magna under DDVP exposure. Chemosphere 120:252–257.

5. Evgenidou E, Konstantinou I, Fytianos K, Albanis T. 2006. Study of the removal of dichlorvos and dimethoate in a titanium dioxide mediated photocatalytic process through the examination of intermediates and the reaction mechanism. J Hazard Mater 137:1056–64.

6. Singh BK. 2008. Organophosphorus-degrading bacteria: ecology and industrial applications. Nature Reviews Microbiology 7:156–164.

7. Zhang X-H, Zhang G-S, Zhang Z-H, Xu J-H, Li S-P. 2006. Isolation and Characterization of a Dichlorvos-Degrading Strain DDV-1 of Ochrobactrum sp. Pedosphere 16:64–71.

8. Harman GE, Howell CR, Viterbo A, Chet I, Lorito M. 2004. Trichoderma species - Opportunistic, avirulent plant symbionts. Nature Reviews Microbiology 2:43–56.

9. Zhang GZ, Zhang XJ, Hong-Mei LI, Guo K, Yang HT. 2016. Isolation and Characterization of the Chlorpyrifos-degrading Trichoderma Strains from the Vegetable Soil in Greenhouse. Biotechnology Bulletin.

10. Tang J, Li Y, Fu K, Yuan X, Gao S, Wu Q, Yu C, Shi W, Chen J. 2014. Disruption of hex1 in Trichoderma atroviride leads to loss of Woronin body and decreased tolerance to dichlorvos. Biotechnology Letters 36:751–759.

11. Zhang T, Tang J, Sun J, Yu C, Liu Z, Chen J. 2015. Hex1-related transcriptome of Trichoderma atroviride reveals expression patterns of ABC transporters associated with tolerance to dichlorvos. Biotechnol Lett 37:1421–9.

12. Jacquet P, Daude D, Bzdrenga J, Masson P, Elias M, Chabriere E. 2016. Current and emerging strategies for organophosphate decontamination: special focus on hyperstable enzymes. Environmental Science and Pollution Research 23:8200–8218.

13. Draganov DI, La Du BN. 2004. Pharmacogenetics of paraoxonases: a brief review. Naunyn Schmiedebergs Arch Pharmacol 369:78–88.

14. Draganov DI. 2010. Lactonases with oragnophosphatase activity: Structural and evolutionary perspectives. Chemico-Biological Interactions 187:370–372.

15. Anonymous. !!! INVALID CITATION !!!

16. Sun W, Chen Y, Liu L, Tang J, Chen J, Liu P. 2010. Conidia immobilization of T-DNA inserted Trichoderma atroviride mutant AMT-28 with dichlorvos degradation ability and exploration of biodegradation mechanism. Bioresource Technology 101:9197–9203.

17. Ceron JJ, Tecles F, Tvarijonaviciute A. 2014. Serum paraoxonase 1 (PON1) measurement: an update. Bmc Veterinary Research 10.

18. Li WF, Costa LG, Furlong CE. 1993. Serum paraoxonase status: a major factor in determining resistance to organophosphates. J Toxicol Environ Health 40:337–46.

19. Teiber JF, Draganov DI, La Du BN. 2003. Lactonase and lactonizing activities of human serum paraoxonase (PON1) and rabbit serum PON3. Biochemical Pharmacology 66:887–896.

20. Bustos N, Cruz-Alcalde A, Iriel A, Fernandez Cirelli A, Sans C. 2018. Sunlight and UVC-254 irradiation induced photodegradation of organophosphorus pesticide dichlorvos in aqueous matrices. Sci Total Environ 649:592–600.

21. Costa LG, McDonald BE, Murphy SD, Omenn GS, Richter RJ, Motulsky AG, Furlong CE. 1990. Serum paraoxonase and its influence on paraoxon and chlorpyrifos-oxon toxicity in rats. Toxicology and Applied Pharmacology 103:66–76.

22. von der Wellen J, Bierwisch A, Worek F, Thiermann H, Wille T. 2016. Kinetics of pesticide degradation by human fresh frozen plasma (FFP) in vitro. Toxicology Letters 244:124–128.

23. Aharoni A, Gaidukov L, Yagur S, Toker L, Silman I, Tawfik DS. 2004. Directed evolution of mammalian paraoxonases PON1 and PON3 for bacterial expression and catalytic specialization. Proceedings of the National Academy of Sciences of the United States of America 101:482–487.

24. Suzuki SM, Stevens RC, Richter RJ, Cole TB, Park S, Otto TC, Cerasoli DM, Lenz DE, Furlong CE. 2010. Engineering Human PON1 in an E. coli Expression System, p 37–45. *In* Reddy ST (ed), Paraoxonases in Inflammation, Infection, and Toxicology, vol 660. Springer-Verlag Berlin, Berlin.

25. Furlong CE, Costa LG, Hassett C, Richter RJ, Sundstrom JA, Adler DA, Disteche CM, Omiecinski CJ, Chapline C, Crabb JW, Humbert R. 1993. Human and rabbit paraoxonases: Purification, cloning, sequencing, mapping and role of polymorphism in organophosphate detoxification. Chemico-Biological Interactions 87:35–48.

26. Furlong CE, Cole TB, Jarvik GP, Pettan-Brewer C, Geiss GK, Richter RJ, Shih DM, Tward AD, Lusis AJ, Costa LG. 2005. Role of Paraoxonase (PON1) Status in Pesticide Sensitivity: Genetic and Temporal Determinants. NeuroToxicology 26:651–659.

27. Furlong CE, Suzuki SM, Stevens RC, Marsillach J, Richter RJ, Jarvik GP, Checkoway H, Samii A, Costa LG, Griffith A, Roberts JW, Yearout D, Zabetian CP. 2010. Human PON1, a biomarker of risk of disease and exposure. Chemico-Biological Interactions 187:355–361.

28. Richter RJ, Jarvik GP, Furlong CE. 2009. Paraoxonase 1 (PON1) status and substrate hydrolysis. Toxicology and Applied Pharmacology 235:1–9.

29. Worth J. 2002. Paraoxonase polymorphisms and organophosphates. The Lancet 360:802–803.

30. Kanamori-Kataoka M, Seto Y. 2009. Paraoxonase activity against nerve gases measured by capillary electrophoresis and characterization of human serum paraoxonase (PON1) polymorphism in the coding region (Q192R). Analytical Biochemistry 385:94–100.

31. Hoskin FCG, Walker JE, Dettbarn WD, Wild JR. 1995. Hydrolysis of tetriso by an enzyme derived from pseudomonas-diminuta as a model for the detoxication of o-ethyl s-(2-diisopropylaminoethyl) methylphosphonothiolate(VX). Biochemical Pharmacology 49:711–715.

32. Mabanglo MF, Xiang DF, Bigley AN, Raushel FM. 2016. Structure of a Novel Phosphotriesterase from Sphingobium sp TCM1: A Familiar Binuclear Metal Center Embedded in a Seven-Bladed beta-Propeller Protein Fold. Biochemistry 55:3963–3974.

33. Valiyaveettil M, Alamneh Y, Biggemann L, Soojhawon I, Farag HA, Agrawal P, Doctor BP, Nambiar MP. 2011. In vitro efficacy of paraoxonase 1 from multiple sources against various organophosphates. Toxicology in Vitro 25:905–913.

34. Wang NN, Yuan L, Dai H, Han ZK, Zhao M. 2011. Effect of PON1 on dichlorvos toxicokinetics. Emergency Medicine Journal Emj 28:313–315.

35. Furlong CE, Cole TB, Walter BJ, Shih DM, Tward A, Lusis AJ, Timchalk C, Richter RJ, Costa LG. 2005. Paraoxonase 1 (PON1) status and risk of insecticide exposure. Journal of Biochemical and Molecular Toxicology 19:182–183.

36. Ruggerone B, Bonelli F, Nocera I, Paltrinieri S, Giordano A, Sgorbini M. 2018. Validation of a paraoxon-based method for measurement of paraoxonase (PON-1) activity and establishment of RIs in horses. Veterinary Clinical Pathology 47:69–77.

37. Kunitake E, Tani S, Sumitani J-i, Kawaguchi T. 2011. Agrobacterium tumefaciens-mediated transformation of Aspergillus aculeatus for insertional mutagenesis. Amb Express 1.

38. Maqbool Z, Hussain S, Imran M, Mahmood F, Shahzad T, Ahmed Z, Azeem F, Muzammil S. 2016. Perspectives of using fungi as bioresource for bioremediation of pesticides in the environment: a critical review. Environ Sci Pollut Res Int 23:16904–25.

39. Collado IG, Aleu J, Pinedo-Rivilla C. 2009. Pollutants Biodegradation by Fungi. Current Organic Chemistry 13:-.

40. Kaushik G, Thakur IS. 2009. Isolation of fungi and optimization of process parameters for decolorization of distillery mill effluent. World Journal of Microbiology & Biotechnology 25:955–964.

41. Kaushik G, Gopal M, Thakur IS. 2010. Evaluation of performance and community dynamics of microorganisms during treatment of distillery spent wash in a three stage bioreactor. Bioresource Technology 101:4296–4305.

42. Xiao Q, Hu B, Yu C, Xia L, Jiang Z. 2006. Optimization of a single-drop microextraction procedure for the determination of organophosphorus pesticides in water and fruit juice with gas chromatography-flame photometric detection. Talanta 69:848–855.

43. Thompson JD, Gibson TJ, Plewniak F, Jeanmougin F, Higgins DG. 1997. The CLUSTAL_X windows interface: flexible strategies for multiple sequence alignment aided by quality analysis tools. Nucleic Acids Research 25:4876–4882.

44. Pfaffl MW, Hageleit M. 2001. Validities of mRNA quantification using recombinant RNA and recombinant DNA external calibration curves in real-time RT-PCR. Biotechnology Letters 23:275–282.

45. Fu K, Fan L, Li Y, Gao S, Chen J. 2012. Tmac1, a transcription factor which regulated high affinity copper transport in Trichoderma reesei. Microbiol Res 167:536–43.

46. Agbagwa IO, Datta S, Patil PG, Singh P, Nadarajan N. 2012. A protocol for high-quality genomic DNA extraction from legumes. Genetics and Molecular Research 11:4632–4639.

47. Yu C, Fan L, Wu Q, Fu K, Gao S, Wang M, Gao J, Li Y, Chen J. 2014. Biological role of Trichoderma harzianum-derived platelet-activating factor acetylhydrolase (PAF-AH) on stress response and antagonism. PLoS One 9:e100367.

48. Nakayashiki H, Hanada S, Quoc NB, Kadotani N, Tosa Y, Mayama S. 2005. RNA silencing as a tool for exploring gene function in ascomycete fungi. Fungal Genetics and Biology 42:275–283.

49. Amitai G, Gaidukov L, Adani R, Yishay S, Yacov G, Kushnir M, Teitlboim S, Lindenbaum M, Bel P, Khersonsky O, Tawfik DS, Meshulam H. 2006. Enhanced stereoselective hydrolysis of toxic organophosphates by directly evolved variants of mammalian serum paraoxonase. Febs Journal 273:1906–1919.

50. Bhattacharyya T, Nicholls SJ, Topol EJ, Zhang RL, Yang X, Schmitt D, Fu XM, Shao MY, Brennan DM, Ellis SG, Brennan ML, Allayee H, Lusis AJ, Hazen SL. 2008. Relationship of paraoxonase 1 (PON1) gene polymorphisms and functional activity with systemic oxidative stress and cardiovascular risk. Jama-Journal of the American Medical Association 299:1265–1276.

